# Periodically driven force protein translocation through a *α*-hemolysin biological nano-pore

**DOI:** 10.1101/462614

**Authors:** M. A. Shahzad

## Abstract

We study the translocation of protein pulled under the action of time periodically external driving force through *α*-hemolysin nano-pore using Langevin molecular dynamical simulation. We observe that time depended external pulling force could enhance to more efficient translocation process as compared to protein translocation driven by constant external pulling force. We characterized the time depended force driven translocation mechanism by studying the gain in translocation as a function of frequency. We also present Golestanian plot which shows the modulated evolutions of number of translocation peptides, and of the probability distribution function with frequency as a results of the transmission of force oscillation to translocation dynamics.

## Introduction

The passage of a molecule through a biological pore, the so-called translocation, is a fundamental process occurring in a variety of biological processes [1], with example such as DNA and RNA transport through a nuclear pore complex, protein transport through membrane channels, and virus injected into cells. The study of the transport of biomolecules through a nanopore is also important for technological application, such as drug delivery and DNA sequencing. In recent past, some theoretical and experimental work has been done where the width of the pore changes during the translocation process. In such translocation process, the dynamical nature of the pore enable the polymer chain [2], and protein [3] to translocate more efficiently as compare to the translocation of biomolecule in static pore. On the experimental side, it has been shown that the translocation of DNA through a nanochannel can be modulated by dynamically changing the cross section of an elastomeric nanochannel device by applying mechanical stress. Time-dependent driving force are also used in the simple unstructured polymer chain translocation process [6–10]. Moreover, using an alternating electric field (time-dependent driving force) in the nanopore has been implied as a source for DNA sequencing [11]. P. Fanzio, et al., [12] has shown that the DNA translocation process through an elastomeric nanochannel device can be altered by dynamically changing its cross section. It has been demonstrated that it is possible to control and reversibly tune the direction of a nano-channel fabricated using elastomeric materials, so to fit the target molecule dimensions. The opportunity to dynamically control the nano-channel dimension open up new possibilities to understand the interactions between bio-molecules and nano-channels, such as the dependence of the translocation dynamics on the channel size, and the effects of moving walls [2]. Here we used a coarse-grained computational model to simulate numerically the translocation of protein through *α*-hemolysin biological pore with time-dependent external pulling force [13–16]. We observed that the periodic pulling force fundamentally changes the behaviors of the translocation time with respect to the angular frequency of the sinusoidal force.

## Model

We used a coarse-grained computational model to simulate numerically the translocation process for the protein through a channel. In such numerical simulation model, the total potential acting on all the residues of the proteins is given by

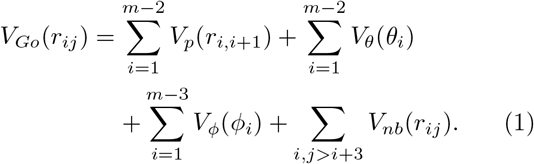

where 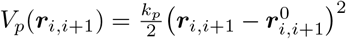 is bond potential with empirical constant 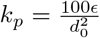 is in term of the equilibrium length parameter *d*_0_, and is in the unit of energy parameter *ϵ*. In our simulation, *d*_0_ = 3.8 Å is the average distance between two adjacent amino acids and *ϵ* sets the energy scale of the model. The angular potential is 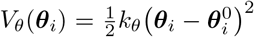 with *k*_θ_ = 20*ϵ* rad^−2^.The dihedral potential are expressed as a function of the twisted angles *ϕ_i_* and 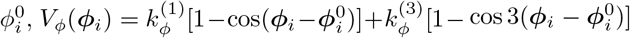, where 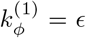 and 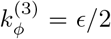 are dihedral constants expressed in term of energy unit *ϵ*. The non-bonded potential *V_nb_* representing the possible long range interaction are express as

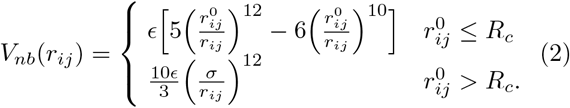

The expression of the pore potential is 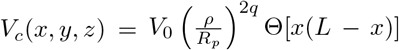, where *V*_0_ = 2*ϵ* and Θ(*s*) = [1 + tanh(*αs*)]/2 is a smooth step-like function limiting the action of the pore potential in the effective region [0, *L*]. *L* and *R_p_* are pore length and radius respectively. Also, 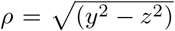 is the radial coordinate. The parameter *q* tunes the potential (soft-wall) stiffness, and *α* modulates the soft step-like profile in the *x*-direction; the larger the *α*, the steeper the step. In our simulation, we consider *q* = 1 and *α* = 2 Å^2^. The driving external force **F**_*ext*_ = *F*_0_[1 + *f* (*t*)]*x̂* acts only in the region in front of the pore mouth *x* ∈ [–2, 0], and inside the channel [0*, L*]. We consider a sinusoidal force given by *f* (*t*) = *F_A_* sin(*ωt* +′*phi*) where *F_A_* is the amplitude, *ω* is the angular frequency and *ϕ* is a constant phase. Pore length *L* = 100 Å and radius *R_p_* = 10 Å are taken from *α*HL structure data. The equation of motion which governs the motion of material point is the well-know Langevin equation which is usually employed in coarse-grained molecular dynamics simulation. Consequently, numerical investigations are performed at constant temperature. The over-damped limit is actually implemented (*r̈* = 0) and a standard Verlet algorithm is used as numerical scheme for time integration [17]. The Langevin equation is 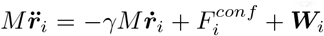 where 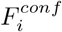 is the sum of all the internal and external forces acting on residue *i*. Here *γ* is the friction coefficient used to keep the temperature constant (also referred as Langevin thermostat). The random force ***W***_*i*_ accounts for thermal fluctuation, being a delta-correlated stationary and standard Gaussian process (white noise) with variance 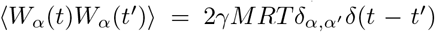.The random force satisfies the fluctuation-dissipation theory; the mean-square of *W* is proportional to the corresponding friction coefficient *γ*.

## Results

We used Langevin molecular dynamics simulation (Verlet integration scheme) to transport protein across a *α*-hemolysin pore using a periodic pulling force. The folding temperature occurred at *T\** = 0.77 in reduced temperature units *R*/ϵ, corresponding to the experimental denaturation temperature *T* = 338 K. This defined the energy scale to the value *ϵ* ≃ 0.88 kcal mol^*−*1^. Using the time scale *t_u_* = *σ*(*M/*120*ϵ*)^1/2^ we can obtained the physical time unit from the simulation time. With *ϵ* ≃ 0.88 kcal mol^−1^, *σ* = 4.5 Å, and assuming the average mass of amino acid is *M* ~ 136 Da, we get *t_u_* ~ 0.25*ps*. In computer simulation the time step and friction coefficient used in the equation of motion (Langevin dynamics) are *h* = 0.001*t_u_* and 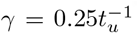, respectively. Further, the unit of force is defined as *f_u_* = *ϵ* Å^−1^ ~ 6 pN. The *f* (*t*) allows the external pulling force to oscillate harmonically, with frequency *ω* and amplitude *A*, about an average width *F*_0_. This defines the *F_min_* = *F*_0_(1 − *F_A_*) and *F_max_* = *F*_0_(1 + *F_A_*), the minimal and maximal of external driving force, respectively. In the simulation, the *F*_0_ = 3 and *F_A_* = 0.9, so that the pore oscillates between *F_min_* = 0.3 and *F_max_* = 5.7.

We used a small protein, named Lipid binding protein (LBP), to understand the basic phenomenology of protein translocation through nanopore. Figure (1) describe the LBP translocation through *α*-hemolysin biological pore pulled with external periodic force. Structurally, LBP is a globular protein of 127 residues with majority of *β*-sheets. Due to its small size, we observed that a pore of radius *R_p_* = 10 Å is enough large that it translocate in hairpin conformation.

**FIG. 1.**
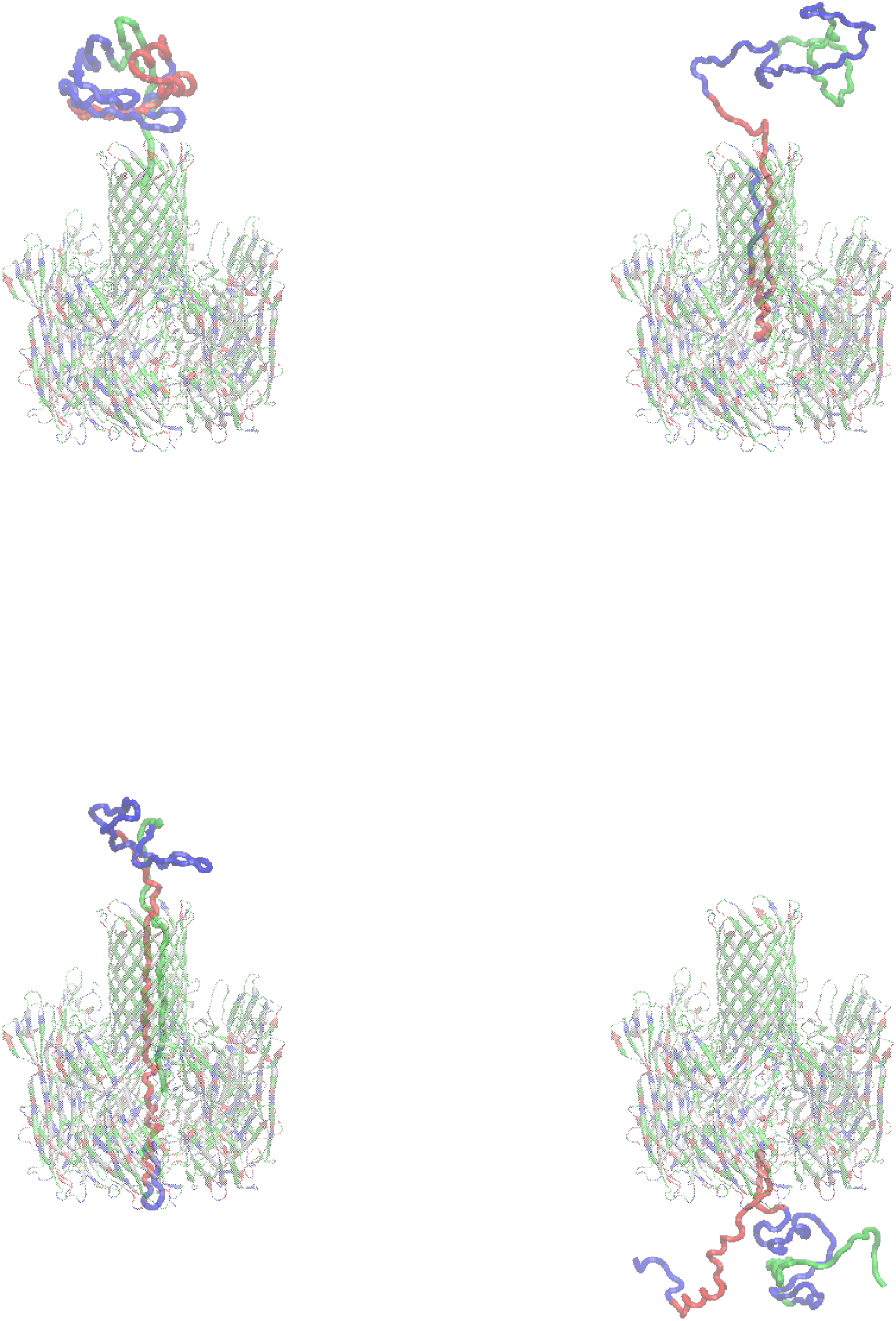
Lipid binding protein transl-location through *α*-hemolysin pore. Upper Panel Left; Initially the folded protein is placed at the CIS-side of the pore. Upper Panel right and Lower Panel Left; Unfold and translocation of protein. Lower Panel Right; Successful translocation and refold of LBP at the TRANS-side.

Figure (2) describes the gain in translocation time defined as *ζ* = *τ*_0_/*τ* as a function of the dimensionless frequency *ω̃* = *ωτ*_0_, where *τ*_0_ is the translocation time in static mode, of the oscillating frequency at phase *θ* = 0. The translocation time displays three different translocation regimes with respect to the frequency of fluctuating external pulling force. In the low frequency limit (*ω̃*< 10^1^), the period *T* of oscillating force is very long and the average translocation time increases steadily until it reached the global maxima. This is because the period of oscillation of the pulling force during the first half cycle is always greater than the average force *F*_0_ of the static mode, hence the protein feels this half cycle before the completion of the translocation process. During the second half cycle the fluctuating force is always smaller than the force of static mode, as a result the translocation time start decreasing. In the limit when *ω̃* → 0, the translocation time approach the time of static pulling force *τ*_0_

**FIG. 2.**
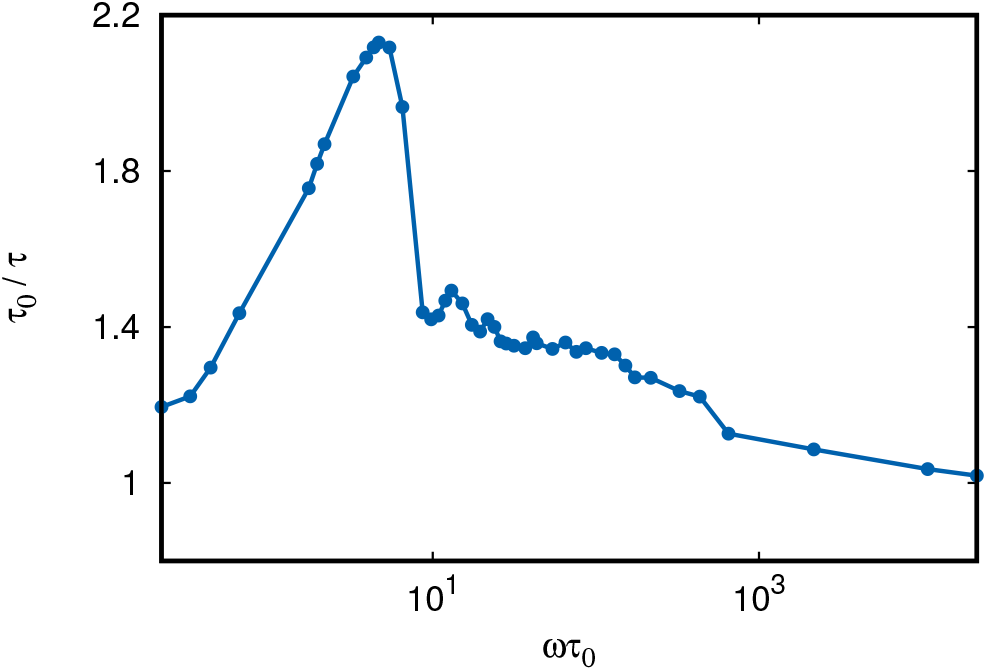
Average translocation time gain *ζ* = *τ*_0_/*τ* as a function of the dimensionless frequency *ω̃* = *ωτ*_0_ with *F*_0_ = 3 and *F_A_* = 0.9. The plot display global maxima follow by series of local maxima and minima.

In the high frequency domain (*ω̃* > 10^3^), a saturation of the translocation time is obtained. The translocation time becomes insensitive at high frequency. In this fast oscillating force regime, the average translocation time become *τ* ~ *τ*_0_, where *τ*_0_ is the translocation time in static mode. Due to high force oscillation rate, the time dependent external force **F**_*ext*_ oscillates many times between *F_min_* and *F_max_* during the course of the translocation process and is averaged out to zero over the whole process. Therefore, the time *τ* ~ *τ*_0_ for *ω̃* → ∞

In the intermediate regime (10^1^ < *ω̃* < 10^3^), the fluctuation of time-dependent radius primary effects the translocation process. We note that the average translocation time possesses one global maxima followed by a series of local minima and maxima. The global maxima of translocation time is obtained at the resonance frequency *ω̃* = 4.6, the so called resonant activation regime. The phenomenon of resonant activation was studied in different problems such as Brownian particle escape dynamics [18], and translocation of polymer in driven systems [2, 6, 7, 19]. The resonance occurred when the polymer driven by a periodic field oscillating at a period comparable with the characteristic time of the crossing dynamics, hence the translocation process accelerates during this period of oscillation. This result is also in agreement with [2], where under different scenario they translocate the polymer through a flickering pore.

The translocation of a protein through a pore driven by oscillating force is further characterized by measuring the average translocation time as a function of the importing force *F* at frequency *ω̃* = 0.0 and *ω̃* = 5.4 with amplitude *A* = 0.9, as shown in Fig. (3). The average was performed over independent runs, excluding those in which protein did not cross the membrane channel from the CIS to the TRANS-side within the waiting time *τ_w_* = 10^5^*t_u_*. In this figure, the brown triangle dot curve is the average time for the translocation of protein through a pore at frequency *ω̃* = 5.4 as compared with the translocation time through a pore with static external pulling force (*ω̃* = 0.0, blue circle dotted line). The average translocation time grows with the decrement of external force *F*, indicating that translocation processes is drastically slowed down. Which indicates the presences of a free-energy barrier due to channel-protein interaction, which the molecule has to overcomes to activate its translocation process. The comparison reveals that at the limit of low external force *F*, the translocation of protein through a pore with oscillating force takes short time as compared to its static counterpart. The difference between the curves representing the translocation times are remarkable particularly at the extreme limit of low force. We note that the translocation with periodic force enhances the translocation process as compared to the static pore. The two curves approximately overlap at the limit of high force due to fast translocation process, hence the static and fluctuating external pulling force becomes indistinguishable.

**FIG. 3.**
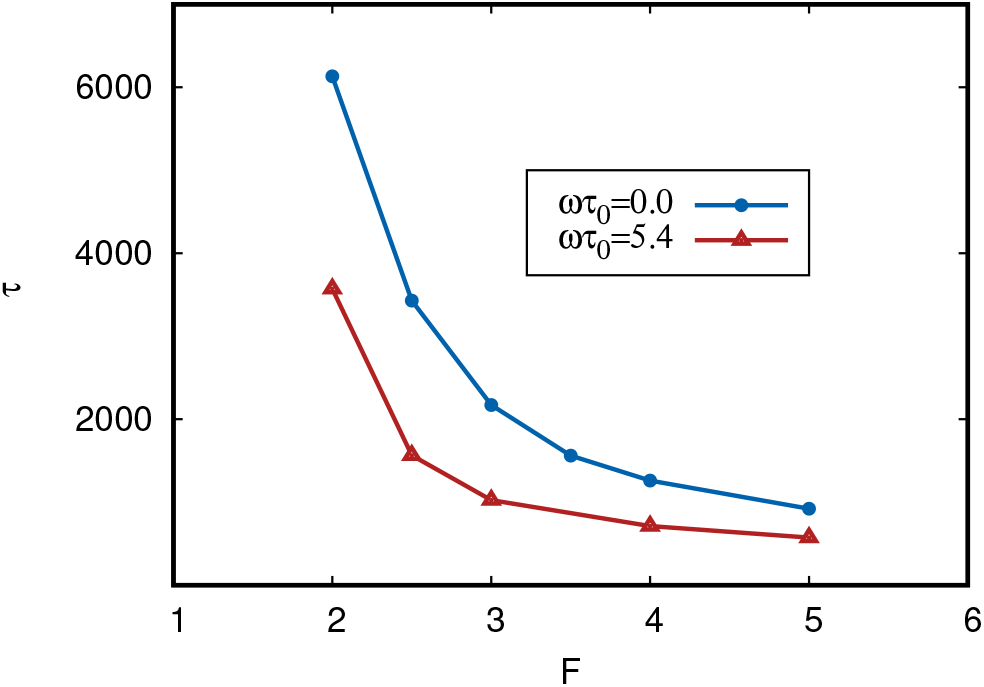
Average translocation time *τ* as a function of homogeneous constant force along *x*-axis *F*_0_ = *F_x_* = *F*.

Figure (4) and (5) shows the distribution of translocation time (the probability density function of first pass time) at two different frequency, *ω̃* = 43 and *ω̃* = 108, respectively (Golestanian plot, [2]). We observed that distribution consists of a series of peaks separated by distinct minima. The series of peaks increase with increase of frequency, as shown in fig. (5) with *ω̃* = 108. The minima of these distinct peaks correspond to the points in the oscillation cycle when the external pulling force is at its minimum, *F_o_*. The detailed dynamics of protein translocation under the action of periodic force is further explore by studding at the number of translocation peptides, *N_trans_*. The *N_trans_* represents the number of peptides that have leave the pore as a function of simulation time. The average modulated dynamics of *N_trans_* with the oscillating force frequency are described in Figure (4) and Figure (5). The comparison of translocation time distribution with the distribution of static mode (black solid line) are also shown.

**FIG. 4.**
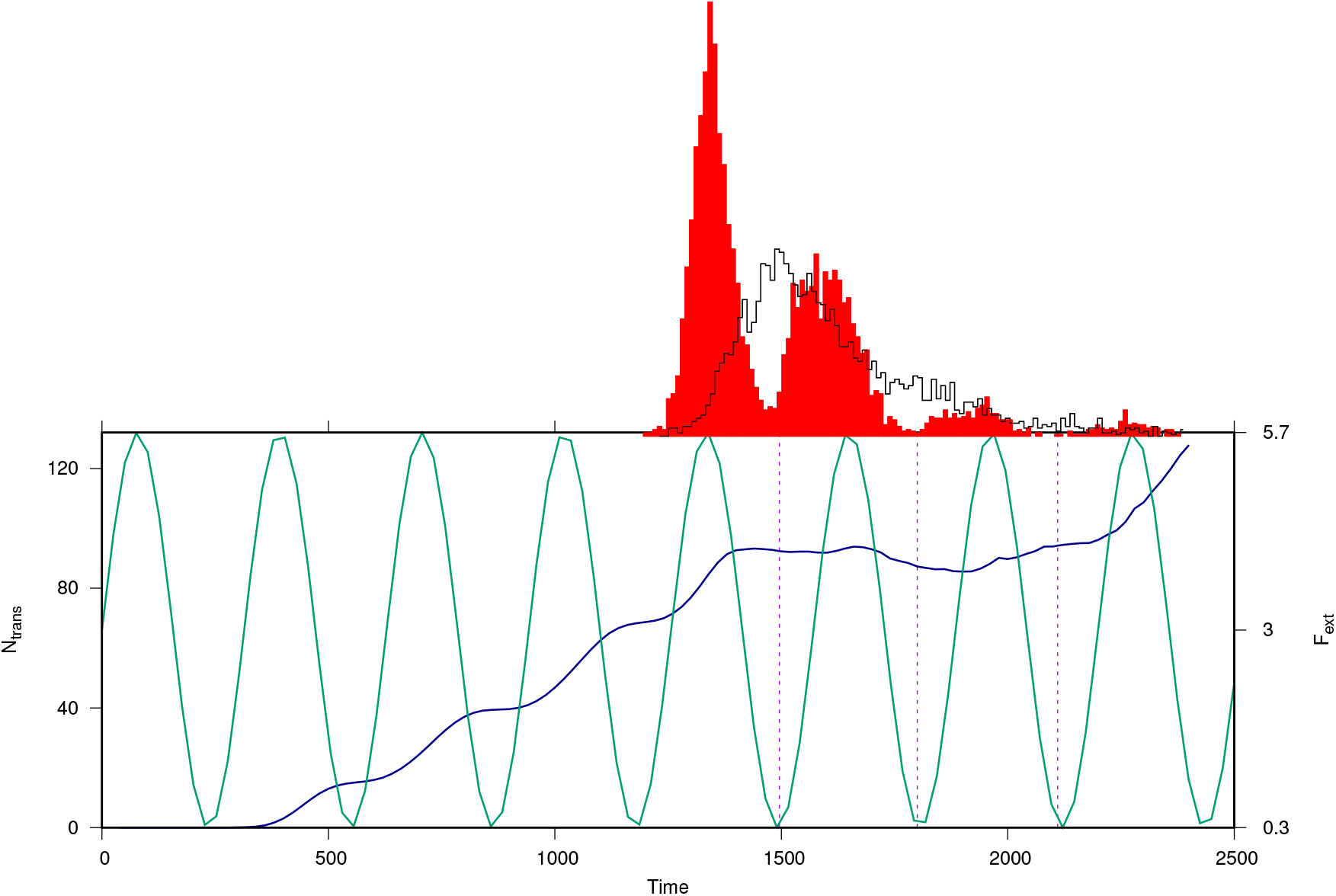
Top Panel:Translocation time distribution as compared with the distribution for the static mode (solid line), and average number of translocation peptides as a function of time (Lower Panel) at *ω̃*= 43 for *F*_0_ = 3 and *F_A_* = 0.9.

**FIG. 5.**
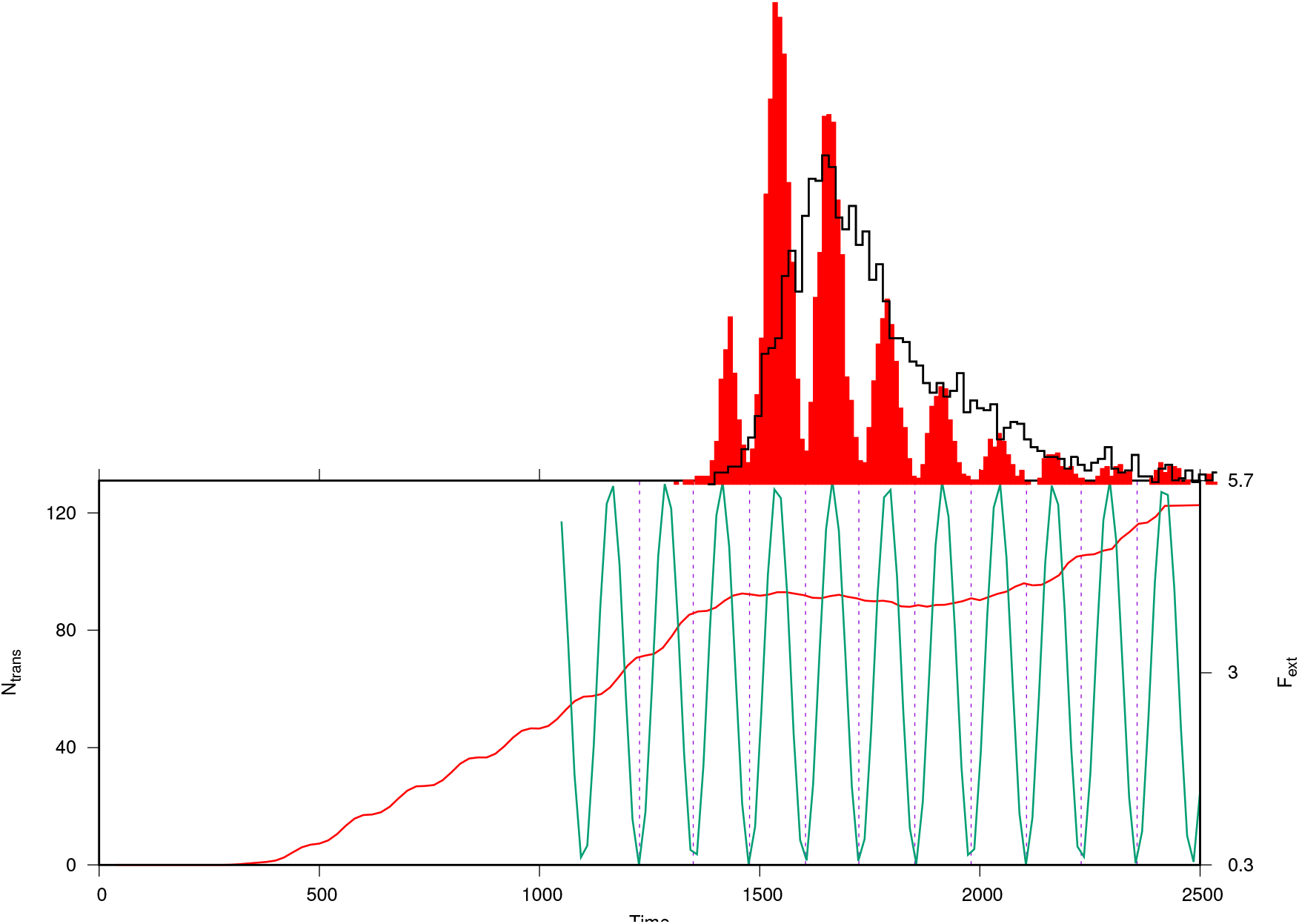
Top Panel:Translocation time distribution as compared with the distribution for the static mode (solid line), and average number of translocation peptides as a function of time (Lower Panel) at *ω̃* = 108 for *F*_0_ = 3 and *F_A_* = 0.9.

## Conclusion

We studied the translocation of proteins through a *α*-hemolysin biological nano-pore using time depended oscillating external pulling force whose force oscillate between the minimal force *F_min_* and the maximal force *F_max_* using Langevin dynamics simulations. In particular, we have extracted the dependence of the average translocation time on the frequency of the oscillating force. The translocation of protein driven by periodic force through a pore play a fundamental role in the dynamics of the system. We can infer from the figure of translocation times as a function of frequency that the translocation process for such a time dependent driven force is sensitive to the frequency of the pulling inward force.

## Supporting Information

Movie S1: This movie file (produced by VMD software) represents the unfold and translocation of LBP through *α*HL nanopore having length *L* = 100 Å and radius *R_p_* = 10 Å.

